# Density-dependent natural selection mediates harvest-induced trait changes

**DOI:** 10.1101/561522

**Authors:** Alix Bouffet-Halle, Jacques Mériguet, David Carmignac, Simon Agostini, Alexis Millot, Samuel Perret, Eric Motard, Beatriz Decenciere, Eric Edeline

## Abstract

Rapid life-history changes caused by size-selective harvesting are often interpreted as a response to direct harvest selection against a large body size. However, similar trait changes may result from a harvest-induced relaxation of natural selection for a large body size via density-dependent selection. Here, we show evidence of such density-dependent selection favouring large-bodied individuals at high population densities, in replicated pond populations of medaka fish. Harvesting, in contrast, selected medaka directly against large-bodied medaka and, in parallel, decreased medaka population densities. Five years of harvesting were enough for harvested and unharvested medaka populations to inherit the classically-predicted trait differences, whereby harvested medaka grew slower and matured earlier than unharvested medaka. We demonstrate that this life-history divergence was not driven by direct harvest selection for a smaller body size in harvested populations, but by density-dependent natural selection for a larger body size in unharvested populations.

## INTRODUCTION

Phenotypic changes caused by harvesting can be exceptionally rapid (Darimont *et al.* 2009) and may have cascading effects on harvesting yields and ecosystem function (Conover & Munch 2002; Dunlop *et al.* 2015). However, the underlying mechanisms that control harvest-induced trait changes are potentially complex and often remain cryptic in empirical studies. The most immediate effect of harvesting is to reduce population density and increase food resources in survivors (Fig. 1, Removal → Density-dependent plasticity pathway), which results in higher rates of somatic growth and reproduction (Verhulst 1838; Hilborn & Walters 1992). However, in parallel, harvesting is often directly size-selective (Fig. 1, Direct harvest selection pathway) and generates complex selective pressures on body size and size-related traits (Matsumura et al. 2011 and references therein).

**Fig. 1.**
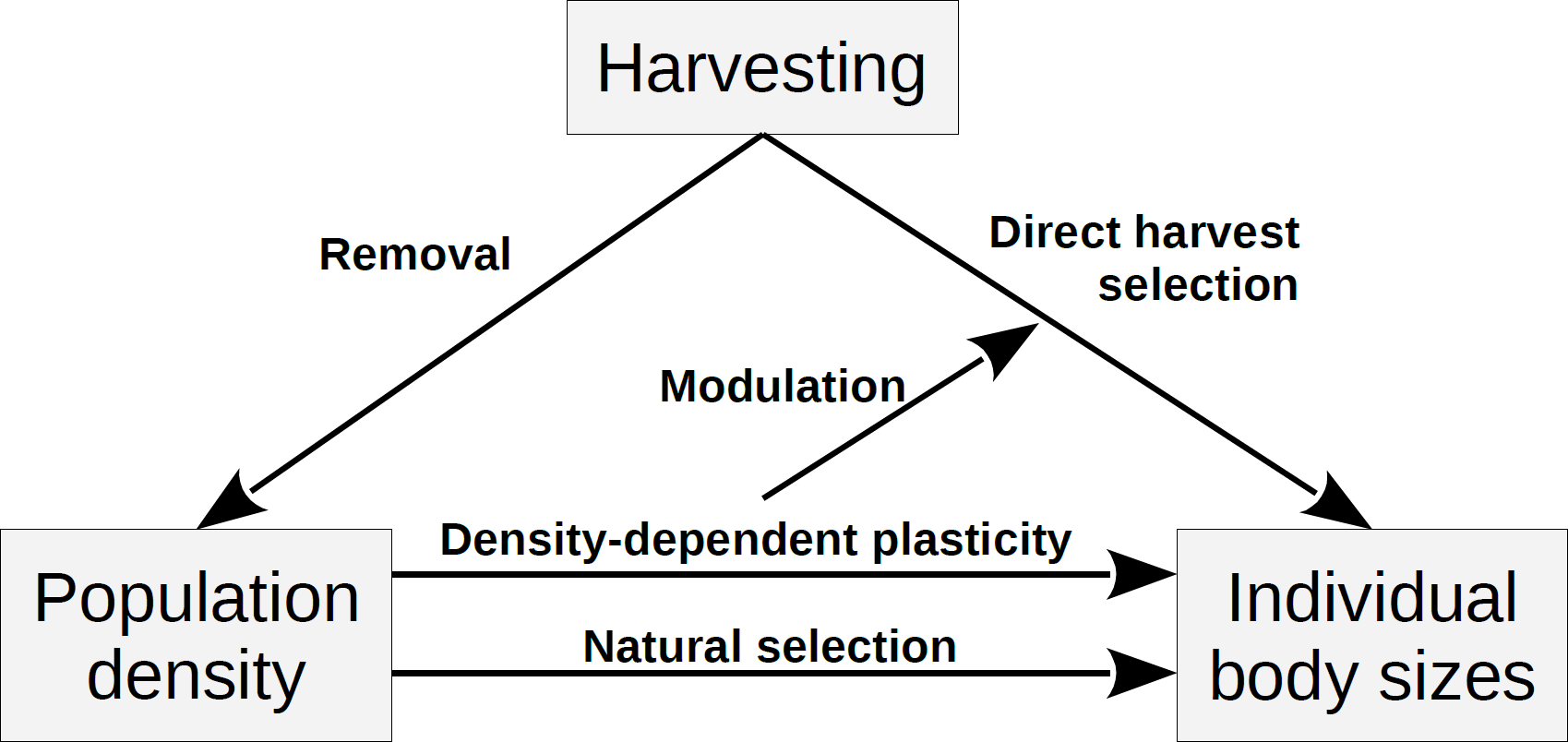
Interaction pathways between size-selective harvesting, population density and individual body sizes. Harvesting simultaneously selects directly against large-bodied individuals (Direct harvest selection pathway) and lowers population density (Removal pathway). The later translates into increased food availability and faster somatic growth in survivors (Density-dependent plasticity pathway). Faster somatic growth rates may shift size-dependent harvesting from targeting both immature and mature fish, to targeting mature only fish (Modulation pathway). Lower population density further relaxes density-dependent natural selection acting on body sizes (Natural selection pathway).

The form and strength of direct harvest selection depends on the specific pattern of selectivity of the fishing gear combined with fishing pressure (Kuparinen *et al.* 2009). For instance, gears targeting large-bodied individuals directly select against fast-growing genotypes (Conover & Munch 2002; Edeline *et al.* 2007, 2009; Swain *et al.* 2007) and, in parallel, select against late-maturing genotypes through reducing life expectancy (Ernande *et al.* 2004; Dunlop *et al.* 2009; Heino *et al.* 2015). Accordingly, a number of empirical and experimental studies have associated harvesting with change towards earlier maturation at a smaller body size and/or towards slower somatic growth (see reviews by Diaz Pauli & Heino 2014; Heino et al. 2015; Kuparinen & Festa-Bianchet 2017). Note, however, that selection for an earlier maturation may also result in evolution of faster somatic growth, allowing for earlier maturation (Dunlop *et al.* 2009; Eikeset *et al.* 2016; Diaz Pauli *et al.* 2017).

Recently, eco-genetic models have further revealed that the presence and strength of density-dependent plasticity in somatic growth can alter the amount and direction of direct harvest selection (Gobin et al. 2018, Modulation pathway in Fig. 1). Such interactions occur because density-dependent plastic changes in somatic growth can shift the timing of maturation in the same direction as harvest selection, thus reducing the strength of direct harvest selection (Lester et al. 2014; Eikeset et al. 2016, but see Arlinghaus et al. 2009). Ultimately, this effect may shift harvesting of large-bodied individuals from removing both immature and mature individuals to removing only mature individuals, in which case selection changes from favouring an early maturation to favouring a late maturation (Ernande *et al.* 2004; Heino *et al.* 2015).

Density-dependent plasticity is not the only pathway through which population density may affect body sizes. The possibility for intraspecific interactions to induce density-dependent, natural-selection on body sizes also exists (Fig. 1, Natural selection arrow). Pioneering studies in *Drosophila* have demonstrated that juvenile (larval) competitive ability in laboratory populations can rapidly evolve in response to crowding, in particular through changes in foraging rates, food conversion efficiency and development time, all of which are traits that affect body size (Mueller 1988, 1997; Sgrò & Partridge 2000; Sarangi *et al.* 2016). More recently, it was shown that density-dependent selection may also be a major driver of trait change in natural populations. For instance, predators relax the strength of density-dependent regulation in wild guppy populations (*Poecilia reticulata*), and degrade the evolved ability of guppies to cope with increasing population density (Bassar *et al.* 2013).

In particular, increased interference competition at high population densities often favours larger-bodied individuals (Post *et al.* 1999). An example of this is the brown anole (Anolis sagrei), where increased interference competition at high population density yields strong natural selection for large body size (Calsbeek & Smith 2007; Calsbeek & Cox 2010). Cannibalism is an extreme form of interference present in a number of taxa, and that also typically selects for large body size (see Claessen et al. 2004 and references therein). In Windermere pike (*Esox lucius*), natural selection is thought to select for larger body sizes through cannibalism (Carlson *et al.* 2007), resulting in a positive relationship between pike density and body size (Edeline *et al.* 2007). Therefore, by lowering population density, harvesting may relax interference competition and cannibalism, in turn decreasing the strength of density-dependent natural selection for large body size. Our aim in this study is to explore this density-mediated, Removal → Natural selection pathway (Fig. 1).

The density-mediated, Removal → Natural selection pathway for harvest selection operates simultaneously with the Direct harvest selection pathway, and both favour smaller body sizes, such that detecting the signature of density-mediated harvest selection is challenging. One way to tackle this challenge is to use an organism in which body size evolves in one direction only. For instance, if body size has already evolved to some lowest possibly physiological limit (Silva *et al.* 2013; Marty *et al.* 2014; Dunlop *et al.* 2015), it can only evolve towards larger body sizes. Therefore, in such an organism any body-size difference between harvested and non-harvested populations would indicate response to density-dependent selection for a larger body size in unharvested populations.

Another potential way to disentangle the effects of natural selection and direct harvest selection is through examining genotype-by-environment interactions on phenotypes. Falconer (1990) showed that, when a trait is selected in environment A, the phenotypic response to selection is expressed in both environment A and other environments (say B), but the amplitude of the phenotypic response to selection is often less in B than in A due to genotype-by-environment interactions. A key environmental effect of harvesting is to increase the levels of available food resources for survivors. Therefore, response to direct harvest selection for a smaller body size should be larger in a high-food than in a low-food environment (Figs. 2a & 2c) while, in turn, response to natural, density-dependent selection for a larger body size should be larger in a low-food than in a high-food environment (Figs. 2b & 2d). Here, we used both approaches to separate the effects of density-mediated and direct harvest selection on body sizes. Specifically, we measured harvest-by-food interactions on somatic growth rate, maturation and energy acquisition rate in the medaka fish (*Oryzias latipes*), an organism that was shown in the laboratory to have an asymmetric body-size evolvability.

**Fig. 2.**
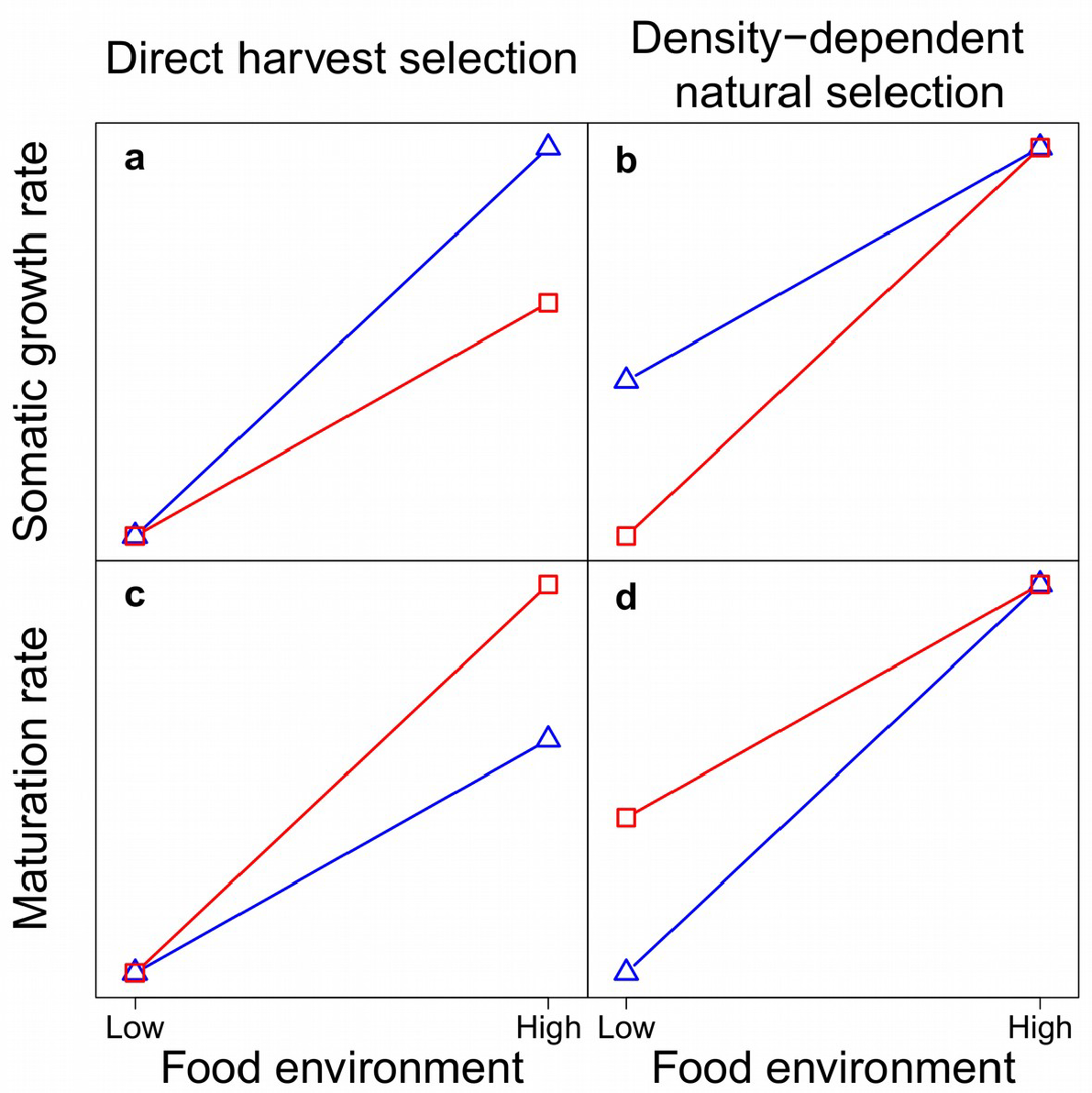
Predicted harvest-by-food interaction patterns on somatic growth rates (e.g., mm day^-1^) and maturation rates (e.g., probability day^-1^) under direct harvest selection *vs*. density-dependent natural selection on body size. Red squares: mean trait value in harvested populations, blue triangles: mean trait value in unharvested populations. Genotype-by-environment interactions often result in response to selection being larger in the environment in which selection was performed than in other environments (Falconer 1990). Harvesting selects for a small body size and simultaneously increases food availability through decreased population density. Hence, body-size response to direct harvest selection is expectedly larger in a high-food than in a low-food environment. In contrast, natural selection for a large body size occurs at high population density and low food. Hence, body-size response to density-dependent natural selection is expectedly larger in a low-food than in a high-food environment.

## MATERIALS AND METHODS

### Pond medaka populations

#### Origin and maintenance

The medaka is an oviparous fish belonging to the group of Beloniformes, a sister group of Cyprinodontiformes which includes killifishes (Kinoshita *et al.* 2009). The medaka, which naturally inhabits slow-moving fresh- and brackish-water habitats of south-east Asia, has a wide thermal range, and can both overwinter under the ice and have a 2-3 months generation time at 27°C in the laboratory. Hence, the medaka is ideal to perform parallel experiments in both the laboratory and in outdoor ponds under temperate latitudes.

Our starting medaka populations descended from 100 parents wild-caught in Kiyosu (Toyohashi, Aichi Prefecture, Japan) in June 2011. Medaka from this populations were shown to respond to selection for a larger body size by evolving faster somatic growth rates and delayed maturation, but were unable to respond to selection for a smaller body size (Le Rouzic *et al.* in press; Renneville *et al.* in press). About 160 progeny were used to seed each of 12 circular outdoor ponds (10 m^2^, 1.2 m deep). Further details on how populations were formed and tanks were installed may be found in the SI Appendix I. No food was added to the ponds so that natural density-dependent processes could take place. To measure the effects of medaka harvesting on their food, we sampled zooplankton (11 dates in 2012) and filamentous algae (7 dates from 2012 to 2013), which are the two major food sources for medaka in ponds (SI Appendix I). Food was manipulated only subsequently during the common garden experiment in the laboratory, so as to test for harvest-by-food interactions on somatic growth, maturation and food intake (see below).

#### Medaka harvesting and phenotyping in ponds

From 2012 to 2016, each of the 12 pond populations was sampled in March before medaka reproduction (pre-recruitment) and in November after medaka reproduction (post-recruitment). Fish were concentrated using a seine net and then fished using handnets (catchability = 98 ± 0.6% SD estimated using removal sampling). All sampled fish were individually weighed to the nearest mg (no anaesthesia required) and estimated for standard body length (from the tip of the snout to the base of the caudal fin) using a body mass-length relationship obtained in March 2013, when fish were also measured individually using the imageJ software (R^2^ = 0.98 on a log-log scale, n = 2722).

Each year in March in the 6 harvested populations, we removed all the fish that were too large to pass through a screen made from 2 mm-spaced parallel bars (i.e., selection on body girth), while in unharvested populations all fish were released after phenotyping. The fishery removed on average 79% of individuals (i.e., 98% catchability × 81% removal rate). Such a high exploitation rate is comparable to those imposed by some industrial marine fisheries (Hutchings 2000; Myers & Worm 2003). In November, all fish from both harvested and unharvested populations were released after phenotyping. The fishing operation and manipulations resulted in an incidental 0.8% mortality rate, which was independent of harvest treatments but that decreased with increasing medaka body size.

#### Larvae counts

In fish, negative density dependence of population dynamics is generally considered to reflect juvenile mortality due to interference competition and cannibalism from large-bodied adults (Ricker 1954; Claessen *et al.* 2004). However, in theory overcompensating recruitment may also operate through decreased adult fecundity. To discriminate between the two mechanisms, we counted the number of newly-hatched larvae hiding in two pairs of floating plastic brushes (summed for analyses) at irregular intervals during the 2014, 2015 and 2016 spawning periods (April to September). In total, for each pond we performed 167 counts spread over 118 dates with one to three counts per date.

### Common garden in the laboratory

Common garden experiments alleviate plastic effects of the environment on the phenotype, and can thus reveal an heritable response to harvesting. We examined harvest-by-food interactions under common garden conditions in the F_1_ medaka generation born from parents sampled in each of the 12 populations. Maternal effects are not alleviated in the F_1_ generation, but they are unlikely to have had large effects in our experiment (see the Discussion).

#### Parental fish

In November 2016, between 6 and 10 medaka (mean 9.6) were randomly sampled from each of the 12 pond populations to serve as parents for a F_1_ generation in the laboratory. These 12 random parental samples, which represented from 3 to 29% of the catch (mean 9%), were maintained in a greenhouse at air temperature in 150L tanks with live food. In January 2017, parental fish were weighed to the nearest mg, measured for standard body length with ImageJ software, and grouped to form 3 breeding pairs per population (36 pairs in total), except for one parental sample which had only one mature female still alive in January 2017 (this female and its mate became parents of all progeny produced from this pond). In parents, we found no significant effect of harvesting on body size (random pond intercept ANOVA, Chisq = 0.353, df = 1, p > 0.552) or body condition (ANCOVA, Chisq = 0.456, df = 1, p > 0.499). Each of the 36 pairs was transferred to the laboratory in a 3.5L aquarium and induced to spawn by progressively raising temperature from 20.0 to 27.0 ± 0.3°C (mean ± SD) and setting a 15-h light : 9-h dark photoperiod. Water conductivity was 375 ± 43 μS/cm.

Dry food (Skretting Gemma Micro, 300 µm pellets) was provided twice per day and live *nauplii* of *Artemia salina* once per day. After initiation of spawning by all breeding pairs, eggs from each breeding pair were collected daily during a 4-day period, enumerated and incubated in separate jars so as to keep track of individual parental identity (but not spawning day). We found no significant effect of harvesting on either probability of a non-zero clutch (zero-inflated negative binomial, random pond intercept, logit scale, estimate of harvest effect = −0.128 (± 0.200 SE), p = 0.522) or on the size of a non-zero clutch (log scale, estimate of harvest effect = 0.071 (± 0.090 SE), p = 0.431).

#### Progeny birth and feeding treatments

We collected larvae hatched from the 7^th^ to the 10^th^ day after the weighted average date of spawning. Larvae born from the same breeding pair on the same day were transferred to 1.5 L aquariums by groups of three larvae (1-4 groups of larvae, mean 2.9, per breeding pair), and were maintained under the same temperature and light regime as their parents. From 15 days post hatch (dph) onwards, we varied resource levels by applying to F_1_ progeny three food environments. In the low-food environment, medaka were fed once every second day with *nauplii* of *Artemia salina*, alternated with dry food. In the high-food environment, medaka were fed twice daily, once with *nauplii* and once with dry food. In the medium-food environment, medaka were fed once daily alternating *nauplii* and dry food. Compared to low-food, the medium- and high-food environments multiplied food supply by 2 and 4, respectively, which loosely corresponds to the *relative* effect of harvesting on zooplankton availability in ponds (see results). Further details on the feeding protocol may be found in the SI Appendix I.

#### Progeny phenotyping

At 15 dph, all F_1_ individuals were weighed and measured as described above and only one individual per aquarium was randomly kept for subsequent phenotyping, making it possible to track individual developmental trajectories.

Individual phenotyping was repeated at 30 dph, 40 dph and then once per week until 90 dph (11 individual measurements for a total of 104 individuals). From 40 dph onwards, phenotyping further included detection of the maturity status from the presence of secondary sexual characters (Yamamoto 1975). Specifically, the maturity criteria were appearance of a round-shaped anal papilla in females, and of the papillar process on the anal fin in males. On average, females matured at 58.4 dph and 15.1 mm, while males matured at 56.5 dph and 14.7 mm. We found no significant difference in somatic growth rate between males and females (random pair intercept ANCOVA, chisq = 0.102, df = 1, p = 0.749).

We measured individual feeding rate three times during a period ranging from 46 to 66 dph. We fasted fish overnight and acclimatized them for five minutes in a 80 mL container under the same temperature and light conditions as during rearing. Fish were then presented with 20 prey (*nauplii* of *Artemia salina*), and we counted the number of prey eaten during 5 minutes.

### Statistical analyses

A full description is given in the SI Appendix I. Briefly, each year in both harvested and unharvested pond medaka populations, the number of age classes was inferred using model-based clustering. The number of fish in each age class at each sampling event was estimated by fitting a mixture of Gaussian distributions to individual body lengths (n = 17908). These estimated numbers allowed us to visualize the strength of negative density-dependence by plotting Ricker stock-recruitment relationships fitted to mean point estimates. We estimated the relationship between individual standard body length and probability to survive through the fishery in March (n = 3970 individuals) using a mixed effects Bernoulli GLM with a logit link function. Finally, medaka larvae and zooplankton counts were modelled using mixed-effects zero-inflated negative binomial models, while % of pond surface covered by filamentous algae was modelled using a negative binomial model.

In the common garden experiment, we estimated harvest-by-food interactions on somatic growth rates and growth trajectories of the F_1_ medaka progeny using a second order polynomial regression of standard body length on age. We modelled medaka maturation using two complementary approaches: probabilistic maturation reaction norms (PMRNs “direct estimation” method, Heino & Dieckmann 2008) and maturation rates (Van Dooren *et al.* 2005; Harney *et al.* 2013). We modelled PMRNs using a Bernoulli GLM that accounted for the effects of both age (days post hatch) and body size (mm) on maturation probability, but that did not include any harvest-by-food interaction so as to gain statistical power in estimating the effect of harvesting on the PMRNs. By doing so, we assumed that the plastic effects of the food environment on the PMRN was fully mediated by somatic growth rates. We modelled harvest-by-food interactions on medaka maturation rates, measured in logit maturation probability day^-1^, using a GLM approximation of a maturation rate model, as described by Harney et al. (2013). Specifically, we fitted to the maturation data a Bernoulli GLM including the time interval between two observations as an offset term. Finally, in the feeding trials, counts of the number of *nauplii* larvae eaten by individual medaka were modelled using a mixed-effects zero-inflated negative binomial model.

## RESULTS

### Age structure and population dynamics in ponds

Both harvested and unharvested pond medaka populations included two age groups corresponding to age 0+ and age 1+ fish (Table S1, Fig. 3a). The experimental fishery targeted medaka larger than 15 mm in standard body length (Fig. 3a, length at 50% removal probability = 18.7 mm) and removed about 50 % of age-0+ recruits and 100 % of 1+ individuals (Fig. 3a), thus reproducing a typical direct harvest selection pattern, which is predicted to favour slow-growing and early-maturing genotypes.

**Fig. 3.**
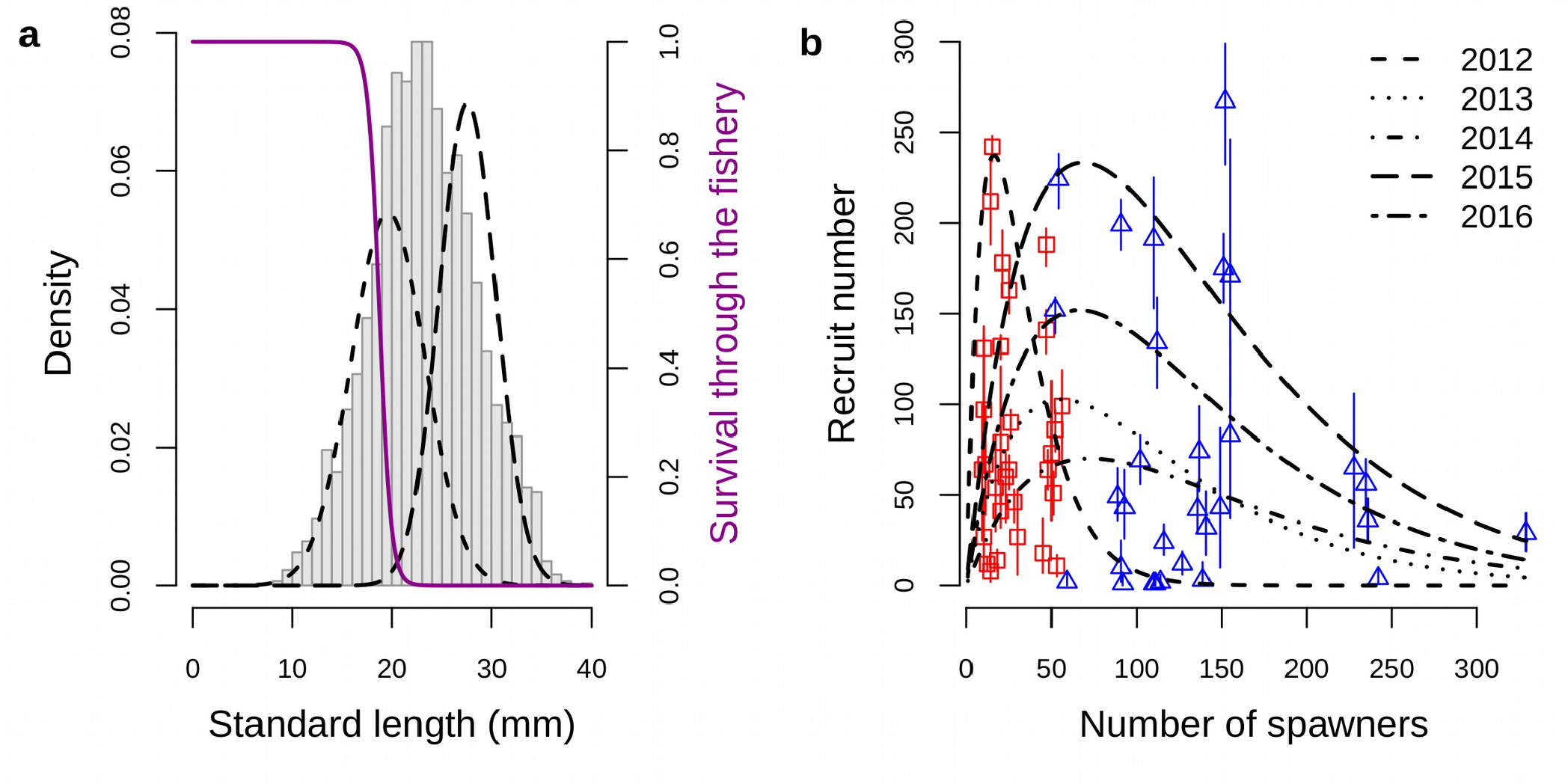
Direct harvest selection and density-dependent natural selection on medaka in ponds. **a: Direct harvest selection on age and size-at-age.** Grey bars represent raw standard length data in harvested populations. Superimposed Gaussians represent mean MCMC estimates for the density of 0+ juveniles (short-dashed curve) and 1+ and older adults (long-dashed curve) individuals. The purple logistic curve is the mean relationship between individual probability to survive through the fishery and standard body length, showing that direct harvest selection was both selecting against an old age and against large-bodied 0+ individuals. **b: Harvesting relaxes negative density-dependence on juvenile medaka recruitment.** Points show mean MCMC estimate for the absolute number of age-0+ recruits with 95% credible intervals from the Gaussian mixture model (see Methods), for harvested (red squares) and unharvested (blue triangles) populations. Each point represents one pond in a given year. Black curves show year-specific Ricker stock-recruitment functions fitted to mean estimates using maximum likelihood.

In parallel with imposing this Direct harvest selection pathway on medaka body size (Fig. 1), our experimental fishery removed negative density-dependence on juvenile recruitment in medaka population dynamics. Pond medaka populations followed “overcompensating” stock-recruitment dynamics (Fig. 3b), which is typical of many other fish populations (Ricker 1954; Hilborn & Walters 1992). Specifically, fishing consistently decreased the stock of spawners (population size in March) below circa 50 individuals (28 on average, red squares in Fig. 2b), a density region in which increasing stock size had a positive effect on the absolute number of age-0+ recruits (black curves, Fig. 3b), indicating demographic “undercompensation” due to density-*in*dependence of vital rates (Bellows 1981). In contrast, unharvested medaka populations had stock sizes above circa 50 individuals (137 on average, blue triangles in Fig. 3b), a density region where increasing stock size had a negative effect on the absolute number of age-0+ recruits, indicating demographic “overcompensation” due to negative density-dependence of vital rates (black curves, Fig. 3b).

Newborn medaka larvae were on average more numerous in unharvested than in harvested populations (P-value = 0.011, Fig. 4, Table S2), i.e., opposite to what one might expect if overcompensating medaka recruitment in unharvested populations occurred via reduced adult fecundity. Hence, as is typical for fish (Ricker 1954), overcompensating recruitment in medaka was instead mediated by increased post-larval (juvenile) mortality, indicating that large-bodied adults dominated smaller-bodied juveniles in pond populations (see also SI Appendix II).

**Fig. 4.**
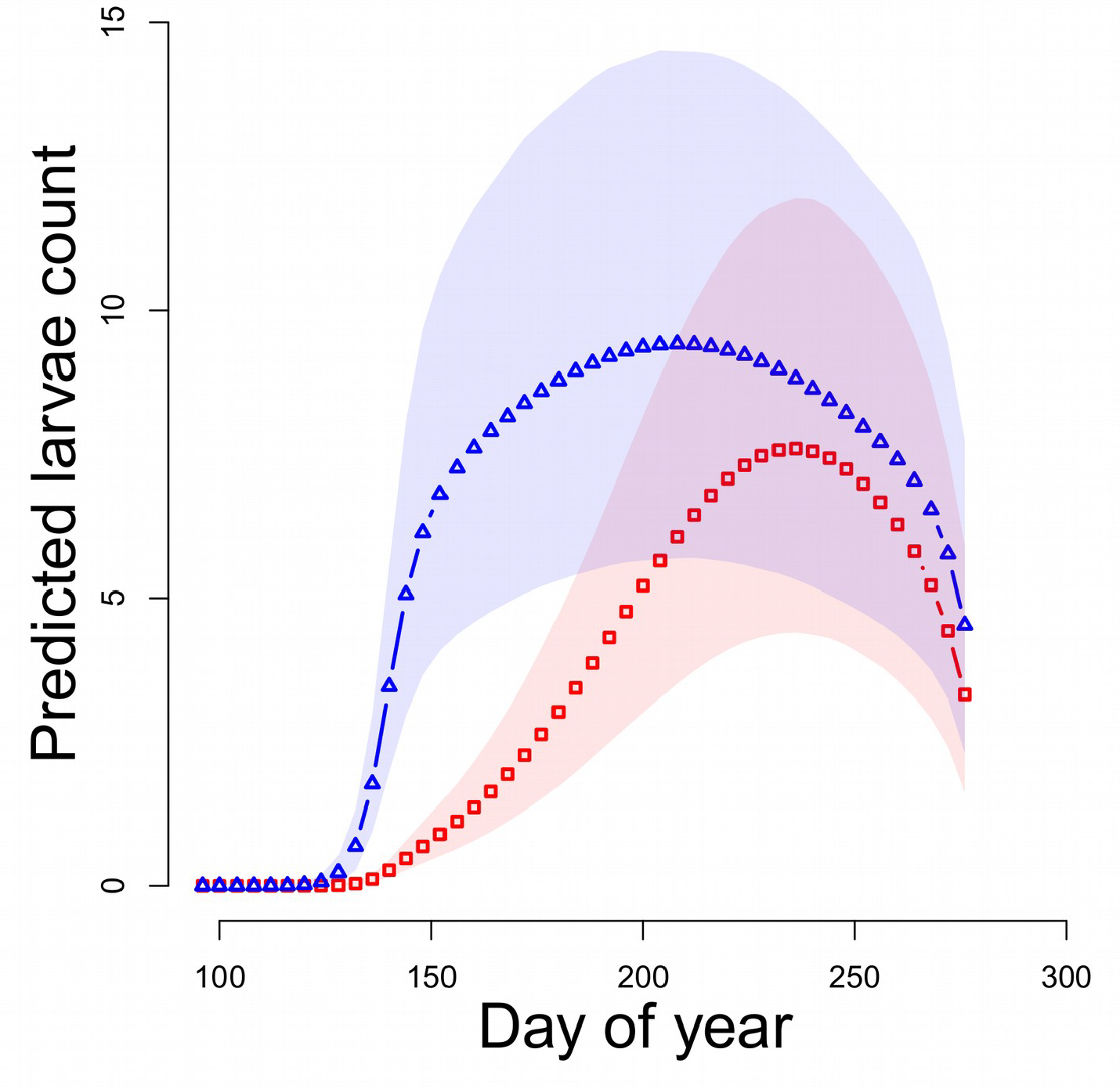
Larvae count seasonal dynamics in ponds during three years. Dots represent mean MCMC estimates for daily counts of newly-hatched larvae for unharvested (blue triangles) and harvested (red squares) populations. Shaded areas show 95% credible intervals around mean MCMC estimates for unharvested (blue) and harvested (red) populations. Raw data are not shown due to a large range (from 0 to 95 larvae, mean 12.4 in unharvested populations; from 0 to 120 larvae, mean 7.4 in harvested populations).

### Food resources in ponds

Fishing for medaka strongly increased abundances of large-bodied zooplankton and filamentous algae, the two major food sources for medaka in ponds. Specifically, medaka fishing multiplied mean abundances of *Asplanchna* sp., Calanoids and Cladocerans by 1.6, 1.9 and 2.9, respectively, and multiplied % pond surface covered by filamentous algae by 14.0 (Table S3).

### Life history in the laboratory common garden

Under all three food environments, F_1_ progeny from harvested populations had significantly lower somatic growth rates than progeny born from unharvested populations (Figs. 5a, MCMC P-values = 0.008 at low food, < 0.001 at medium food and = 0.002 at high food), resulting in a similar effect of harvesting on somatic growth trajectories (Fig. 6a). Accordingly, a deviance analysis shows that there was no significant harvest-by-food interaction on medaka somatic growth rates (P-value of Age × Harvesting × Food interaction = 0.265, Table S4), indicating that the harvest-induced decrease in somatic growth was food-independent. This absence of any harvest-by-food interaction does not permit selective drivers to be disentangled, and therefore can not support any of our two predictions (Figs. 2a, 2b).

**Fig. 5.**
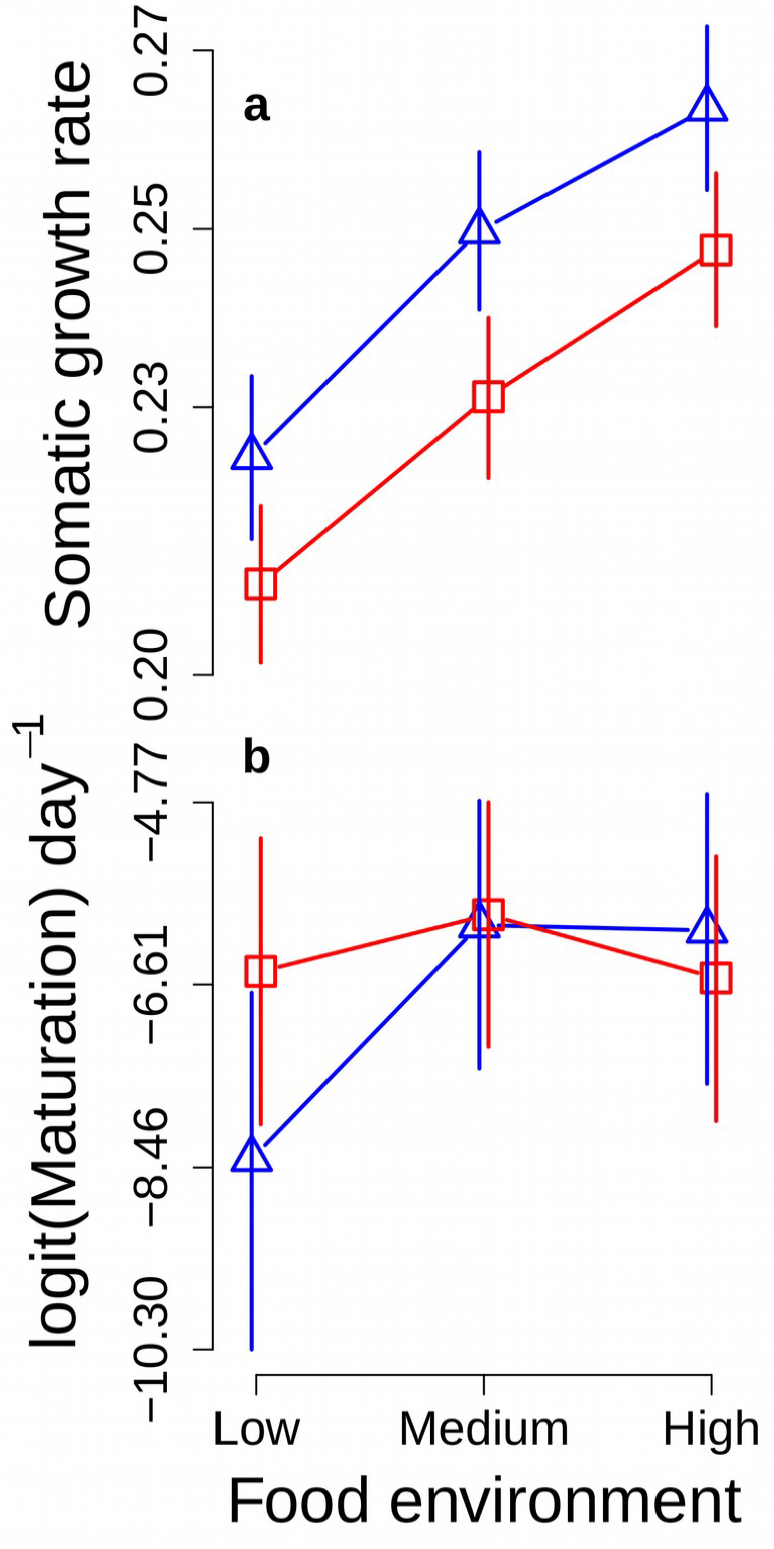
Harvest-by-food interaction patterns on somatic growth rates and maturation rates in individually-raised F_1_ progeny in the laboratory. Points represent mean MCMC estimates with 95% credible intervals for medaka originating from unharvested (blue triangles) and harvested (red squares) populations, and maintained in a low-food, medium-food or high-food environment. Under low food conditions, offspring from harvested populations grew slower and matured at a faster rate than unharvested populations. Somatic growth rates are in mm day^-1^, maturation rates are in logit of maturation probability per day.

**Fig. 6.**
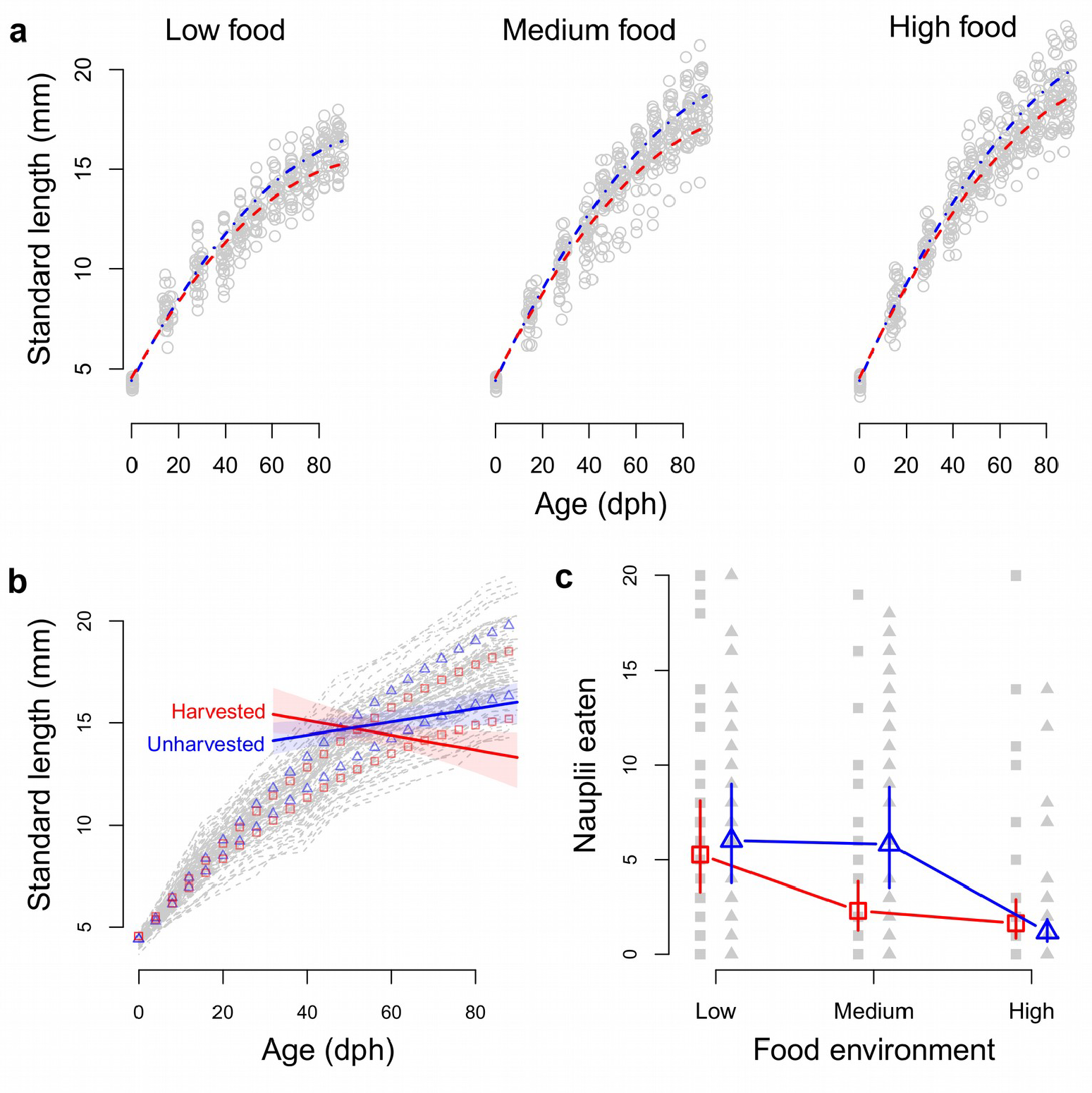
Life-history of individually-raised F_1_ progeny in the laboratory. **a: Somatic growth trajectories.** Somatic growth curves from mean MCMC parameter estimates for individuals originating from unharvested (dot-dashed blue curves) and harvested (dashed red curves) populations in a low-, medium- or high-food environments. Grey dots show the raw data. Dph: days post hatch. **b: Probabilistic maturation reaction norms (PMRNs).** Solid lines show MCMC mean estimates for 50% maturation probability with 95% credible intervals (not to be confounded with the PMRN width). In the background, thin grey curves show raw somatic growth trajectories for medaka originating from unharvested (dot-dashed) and harvested (dashed) populations, and dots show mean somatic growth trajectories from panel (a) for unharvested (blue triangles) and harvested (red squares) medaka in a low-food and high-food environments (medium-food environment ommited for clarity). **c: Feeding behaviour.** Coloured, open points symbols show mean MCMC estimates with 95% credible intervals for the number of prey eaten by medaka originating from unharvested control (blue triangles) and harvested (red squares) populations and maintained in a low-, medium- or high-food environment. Grey, filled symbols show the raw data.

In a low-food environment (MCMC P-value = 0.015), but not in a high-food or medium-food environments (MCMC P-values = 0.881 and 0.506, respectively), maturation rates were higher in F_1_ progeny from harvested populations than from unharvested populations (Fig. 5b). The deviance analysis confirmed the significant Age × Harvesting × Food interaction (P-value < 0.001 in Table S4). This harvest-by-food interaction on maturation rates resulted in harvesting having no influence on the height of medaka PMRN, but on its slope, which shifted from positive to negative (Fig. 6b). Specifically, in a high-food environment medaka reached a 50% maturation probability at around 50 days post hatch (dph) in both harvested and unharvested populations (Fig. 6b). In contrast, in a low-food environment the 50% maturation probability was reached at around 65 dph by harvested medaka, but was reached at about 80 days by unharvested medaka (Fig. 6b). Taken together, these results support the prediction that medaka maturation rates responded to selection in a low-food environment, i.e., responded to density-dependent natural selection (Fig. 2d), but not to direct harvest selection (Fig. 2c). Note that we did not find any strong population pattern on somatic growth rates or maturation probabilities (Fig. S2).

### Feeding trials in the laboratory

F_1_ progeny from harvested and unharvested medaka populations ate a similar number of prey in a low-food and in a high-food environments (Fig. 6c, MCMC P-values = 0.673 and 0.405, respectively), indicating no difference in food acquisition rate. In contrast, in a medium-food environment F_1_ progeny from harvested populations ate significantly less prey than progeny from unharvested populations (Fig. 6c, MCMC p-value = 0.015).

## DISCUSSION

Our results show that, as expected, harvested medaka grew slower and matured earlier than unharvested medaka. Convergent elements suggest that these differences were inconsistent with an adaptive response of medaka owing entirely to direct harvest selection for a smaller body size, but that were driven in part by natural selection for larger body size (Fig. 1).

In ponds, medaka populations included two age classes which, in March of year *t*, were age 0+ fish born in summer of year *t*-1 and age 1+ fish born in summer of year *t*-2. In harvested populations, however, almost all 1+ fish were removed by the fishery in March (Fig. 3a) and only 0+ individuals were effectively able to reproduce, similar to the wild in Japan where medaka die during their first reproductive season (Edeline et al. 2016 and references therein). Therefore, in ponds fishing mortality replaced the natural mortality regime that prevails in the wild, and harvesting is thus unlikely to have driven any large response in medaka life histories. In contrast, in unharvested populations large-bodied individuals were favoured at high population densities (Fig. 3b) and enjoyed a second reproductive season (full discussion in SI Appendix II), probably causing the observed changes towards faster somatic growth and delayed maturation.

This conclusion is supported by harvest-by-food interactions on maturation. Both probabilistic maturation reaction norms and maturation rates consistently showed that harvested medaka matured significantly earlier than unharvested medaka in a low-food environment only. This result suggests that medaka maturation responded to selection in a low-food environment (Falconer 1990) and, hence, responded to selection under high population densities in unharvested populations (Fig. 2). This result fits with results in the Trinidadian killifish (*Rivulus hartii*), in which predation-induced change to earlier maturation was more pronounced in a high-food environment, presumably because predators decrease the population density of *Rivulus* and thus favour genes conferring an early maturation in a high-food environment (Walsh & Reznick 2008, see also Walsh & Reznick 2010, 2011).

Finally, the primacy of natural selection as a driver of life-history divergence among harvested and unharvested pond medaka populations is also supported by a previous selection experiment in the laboratory. In this experiment, medaka were either randomly size-selected (control line), selected against a large body size or selected against a small body size, with populations sizes (about 200 individuals) and intensity of selection on body size (80%) similar as those in the ponds. In the laboratory, response to size-selection was asymmetric: medaka were unable to respond to selection for a smaller body size, but were able to evolve delayed maturation and faster somatic growth in response to selection for a larger body size (Le Rouzic *et al.* in press; Renneville *et al.* in press). This is an important result suggesting that, in ponds also, medaka did not respond to direct harvest selection for a smaller body size but rather to density-dependent natural selection for a larger body size (Fig. 1).

The absence of any harvest-by-food interaction on medaka somatic growth rates did not support either harvest or natural selection exclusive as a primary driver of trait change. This negative result may have two mutually-exclusive causes: (1) harvest selection at high food and natural selection at low food were identical in strengths and, at the same time, induced similar correlated responses in a low- and high-food environments, respectively, or (2) somatic growth rates are not subject to gene-by-food interactions in medaka. Identical strengths of natural and harvest selection are improbable because harvest selection was shown to be stronger than natural selection in multiple systems (Darimont *et al.* 2009), and similar correlated responses in multiple environments are uncommon (Falconer 1990). In contrast, somatic growth rate was reported not to show any gene-by-food interaction in several experiments in mice and rats (Falconer 1990). Hence, we conclude that medaka somatic growth rates were most likely not subject to any gene-by-food interaction, such that selection on somatic growth in one food environment resulted in a similar response in any food environment.

### Rates of food acquisition

In the medaka, it was recently shown in the laboratory that selection for a larger body size favoured higher rates of food acquisition in females and lower boldness in males (Diaz Pauli *et al.* 2019), supporting the prediction that size-dependent selection may alter metabolic rates (Claireaux et al. 2018 and references therein). Combining our results on rates of food acquisition, somatic growth and maturation allows us to propose qualitative hypotheses for how different energy pathways responded to size-dependent selection in different food environments.

In a low-food environment, unharvested and harvested medaka had similar food-acquisition rates (Fig. 6c), but unharvested medaka had higher somatic growth rates (Figs. 5a, 6a) and delayed maturation (Figs. 5b, 6b), suggesting energy re-allocation from reproduction to somatic growth. In a medium-food environment, unharvested medaka had higher food acquisition rates than harvested medaka, which may explain also why unharvested medaka grew faster (Figs. 5a, 6a), but the higher feeding rate of unharvested medaka did not induce any earlier maturation (Figs. 5b, 6b), again suggesting energy reallocation to somatic growth. Finally, in a high-food environment unharvested and harvested medaka had similar rates of food intake (Fig. 6c) and maturation (Figs. 5b, 6b), but unharvested medaka grew faster (Figs. 5a, 6a), suggesting that they had a higher energy assimilation efficiency. Note that our conclusions hold true only if energy pathways other than somatic growth and maturation show negligible response to harvesting.

It is also important to keep in mind that we obtained our results in a simplified experimental system, where medaka could virtually not avoid interacting with the fishing gear (98% catchability) and, hence, where the fishery selected directly on body size only (Fig. 1). In the wild, fish can avoid interacting with the fishing gear through adopting different escape behaviours or habitat choices, and fisheries thus select directly on both behaviour and body size in parallel (Diaz Pauli & Heino 2014; Claireaux et al. 2018). Compared to strict size-dependent selection, this added complexity might potentially lead to different rearrangements in the rates of energy acquisition, assimilation and/or allocation.

### Maternal effects

By examining harvest-by-food interactions in F_1_ medaka progeny, our design avoided the possible selective effects of the captive environment on medaka phenotypes (i.e., domestication), but did not remove possible maternal effects, defined as traits or genes expressed in the mother that influence offspring phenotypes (Lynch & Walsh 2018). Parental body condition was not found to differ between harvested and unharvested populations in either sex, which makes maternal or paternal effects unlikely to have occurred. Additionally, in fish environmental maternal effects often occur when high-quality maternal environments result in larger-sized eggs and in faster early somatic growth in the offspring, an effect that generally vanishes as individuals develop and approach maturity (Einum & Fleming 1999; Heath *et al.* 1999; Lindholm *et al.* 2006). We did not measure egg size in our experiment. However, F_1_ progeny born from females experiencing a high-food environment (i.e., from harvested population) had a similar size at hatch and grew slower, not faster, than did F_1_ progeny born from females experiencing a low-food environment (i.e., from unharvested populations). This result is opposite to what we would expect from an environmental maternal effect on early somatic growth rates. Therefore, we conclude that environmental maternal effects were small and did not strongly influence our results.

## Conclusions

For the first time, we provide experimental evidence suggesting that life-history divergence between harvested and unharvested populations may result from natural, density-dependent selection for a larger body size in unharvested populations. This result strengthens the mounting evidence showing that density-dependent selection may be a primary driver of trait dynamics (Mueller 1988, 1997; Moorcroft *et al.* 1996; Reznick *et al.* 2002; Calsbeek & Smith 2007; Edeline *et al.* 2007; Calsbeek & Cox 2010; Sarangi *et al.* 2016). The effects of density-dependent natural selection, which often favours larger body sizes at higher population densities, come in conflict with the effects of density-dependent plasticity, which favours smaller body sizes at higher densities due to food limitation or social stress (Edeline *et al.* 2010). This conflict blurs the phenotypic effects of density-dependent selection, which are thus likely to remain unnoticed (Wolf 2003; Hadfield *et al.* 2011; Kinnison *et al.* 2015) and, hence, might be more common than the literature suggests and might help to explain why the relationship between biomass productivity and population density is often tenuous (Vert-pre *et al.* 2013). Density-dependent natural selection has important ramifications to the management of harvested populations (Engen *et al.* 2014) and may drive eco-evolutionary feedback loops, which are critical for the maintenance of biodiversity in the context of global changes (Dieckmann & Ferrière 2004; Edeline & Loeuille 2020). In particular, introduction of invasive species, habitat destruction, climate warming and harvesting are all strong disruptors of both population density and size structure and, hence, are potentially strong drivers of density-dependent eco-evolutionary feedbacks on body size.

## Acknowledgements

We are indebted to Prof. Kiyoshi Naruse (NIBB, Okazaki, Japan) for his support in obtaining and maintaining wild medaka from Kiyosu. Tony Masson, Solène Boursault, Angevine Masson, and Nicolas Florès contributed to count medaka larvae in ponds. Iago Bonnici, Stéphane Loisel, Antoine Vallier and Paul Hübner contributed to counting zooplankton, and David Rozen, Gérard Lacroix and Stéphane Loisel contributed to collecting data on filamentous algae. We thank Prof. L. Asbjørn Vøllestad (Universitetet i Oslo) and Nicolas Loeuille (Sorbonne Université) for helpful comments on an early version of the manuscript. Jenilee Gobin (Trent University) and four anonymous reviewers further provided highly relevant comments that helped us to strongly improve the quality of later versions.

## Author contributions

ABH performed the laboratory F_1_ experiment, contributed to data analysis, wrote the first draft of the manuscript and contributed to subsequent versions. EE designed the study, contributed to the pond experiment, performed data analysis, and led manuscript writing from the second version. JM, DC, SA, AM, SP, EM and BD performed the pond and laboratory experiments.

## Data archiving statement

All data used in this paper will be archived.

## Ethical statement

The protocols used in this study were designed to minimize discomfort, distress and pain of animals, and were approved by the Darwin Ethical committee (case file #Ce5/2010/041).

## Supplementary Information

SI Appendix I: Supplementary Methods.

SI Appendix II: Natural selection on body size in medaka.

SI Appendix III: Supplementary Results:

- Table S1: Inference of number of age classes in pond medaka populations.
- Table S2: Structure and MCMC parameter estimates for models 4-6 and 8.
- Table S3: Effect of medaka fishing on medaka food in ponds.
- Table S4: Analysis of deviance.
- Fig. S1: Experimental design.
- Fig. S2: Random effects of breeding pairs on their progeny’s somatic growth rate and maturation probability.

