## Supplementary Material for "Density-dependent natural selection mediates harvest-induced trait changes"

**This PDF file includes:**

SI Appendix I: Supplementary Methods.
SI Appendix II: Natural selection on body size in medaka.
SI Appendix III: Supplementary Results:

- Table S1: Inference of number of age classes in pond medaka populations.
- Table S2: Structure and MCMC parameter estimates for models 4-6 and 8.
- Table S3: Effect of medaka fishing on medaka food in ponds.
- Table S4: Analysis of deviance.
- Fig. S1: Experimental design.
- Fig. S2: Random effects of breeding pairs on their progeny's somatic growth rate and maturation probability.

### APPENDIX I. SUPPLEMENTARY METHODS

#### Origin of the experimental medaka populations

Our start medaka populations descended from 100 parents wild-caught in Kiyosu (Toyohashi, Aichi Prefecture, Japan) in June 2011. These 100 breeders were brought to the Centre de Recherche en Ecologie Expérimentale et Prédictive near Paris, France (CEREEP – ECOTRON Île-de-France, www.cereep.ens.fr), and maintained in groups of 20 individuals in five 20 L aquariums where they mated randomly. Eggs were collected from July to September 2011 and incubated in Petri dishes. Hatched larvae were randomly assigned to 12 circular outdoor ponds (3.57 m diameter, 1.2 m deep) at a density of about 160 larvae per pond which, we assumed, was sufficient to prevent founder effects.

Prior to medaka introduction, the 12 ponds were bottom-coated with a 5 cm layer of Loire River sand, filled with tap water and mildly enriched with a plant fertilizer. After a few weeks of algal development, tanks were seeded with a diverse community of zooplankton collected from surrounding water bodies. Medaka introduction was performed after ponds had reached a clear-water state indicating algal control by zooplankton. After introduction, two pairs of floating plastic brushes were placed in each tank to provide fish with a spawning substrate and shelter for larvae. Afterwards, ponds received only rain water and aerial deposits. Each pond was covered with a net to prevent avian predation, and outlets were secured with a stainless steel filter to prevent any fish or egg escapement.

**Monitoring medaka food in ponds**

In 2012, we measured the effect of medaka fishing in March on zooplankton and filamentous algae, which are primary food sources for medaka in ponds, during the following months. In each pond on 11 dates (April $11^{th}$ and $27^{th}$, May $9^{th}$ and $23^{rd}$, June $6^{th}$ and $13^{th}$, July $4^{th}$ and $18^{th}$, August $22^{nd}$ and September $18^{th}$), zooplankton was sampled from twelve, 2 L water-column samples homogeneously spread across the pond. The resultant 24 L were filtered on a 50 µm filter and the retained zooplankton were fixed in 99% ethanol before subsequent enumeration of rotifers (*Asplanchna* sp. *vs*. other rotifers), Copepod *nauplii*, copepodite stages of calanoid and cyclopid Copepods, and Cladocerans using either a binocular microscope, the ZooScan (Gorsky *et al.* 2010) or the FlowCam (Sieracki *et al.* 1998). Percentage of pond surface covered by filamentous algae was visually estimated by multiple observers on 5 dates in 2012 (May $9^{th}$, September $18^{th}$ and $24^{th}$, October $9^{th}$ and $23^{rd}$) and 2 dates in 2013 (May $22^{nd}$ and July $12^{th}$).

**Feeding of $F_1$ progeny in the laboratory**

In the low-food environment, medaka were fed once every second day with 2 mL of a solution containing *nauplii* of *Artemia salina* (INVE Aquaculture SEP-Art cysts) providing 8.2 ± 0.7 (mean ± SD) mg dry weight of *nauplii* (assuming a 40% dry weight yield from cysts from INVE technical sheets), alternated with dry food (Skretting Gemma Micro, see below). In the high-food environment, medaka were fed twice daily, once with *nauplii* and once with dry food. In the medium-food environment, medaka were fed once daily alternating *nauplii* and dry food.

Dry food doses (measured with volumetric spoons) and pellet sizes were adjusted during fish development to fit with the ontogenetic increase in energy needs and prey size. We computed theoretical daily needs in dry food mass per fish as $M(a)\times 0.30\left(M(a)/M_b\right)^{-0.25}$, where $M(a)$ is individual fish body mass at age *a* (as estimated from previous laboratory data on mass-age relationship in Renneville et al. 2020), $M_b$ is individual fish body mass at birth, and the -0.25 exponent follows from the metabolic theory of ecology (Brown *et al.* 2004). To roughly follow these theoretical needs, medaka received daily 2, 3 and 7 mg of dry food from ages 0 to 40, 40 to 60, and 60 days-post-hatch (dph) onwards, respectively. Pellet size (µm) was 100% 150, 50% mixture of 150-300 and 100% 300 from ages 0 to 20, 20 to 40, and 40 dph onwards, respectively.

**Statistical analyses**

*Fishery selection in ponds*

We estimated the relationship between individual standard body length and probability to survive through the fishery using a Bernoulli GLMM with a logit link function:

$$\begin{aligned} & y_i \sim B(p_i) \\ & \ln\left(\frac{p_i}{1-pi}\right)=\alpha_0+\alpha_{k[i]}+\left(\beta_0+\beta_{k[i]}\right) Sdl_i \\ & \begin{pmatrix}\alpha_k \\ \beta_k\end{pmatrix} \sim N\left(\begin{pmatrix}0 \\ 0\end{pmatrix}, \begin{pmatrix}\sigma_\alpha & \rho\,\sigma_\alpha\sigma_\beta \\ \rho\,\sigma_\beta\sigma_\alpha & \sigma_\beta\end{pmatrix}\right)\end{aligned} \qquad (2),$$

where $B$ is the Bernoulli distribution, subscripts $i$ and $k$ index individuals (*n* = 3970) and sampling events, respectively, to which individuals belong, and ln is the natural logarithm. There were $n$ = 6 fished populations and $n$ = 5 sampling years, yielding $k = 1,2 \ldots 30$ sampling events. Eq. 2 indicates that we modelled the intercept and slope of the survival-length relationship as normally-varying among sampling events $k$, including a correlation parameter $\rho$ between intercept and slope. Parameter estimates $\alpha_0$ and $\beta_0$ from Eq. 2 define a mean size-dependent survival function as plotted in Fig. 3a in the main text.

*Medaka aging and population dynamics in ponds*

Medaka juveniles are too small to be tagged and, unlike in Japan (Terao 1985; Edeline *et al.* 2016), no winter check was deposited in medaka otoliths in our experimental populations. We therefore relied on an analysis of body length-frequency distributions to infer medaka age using model-based clustering in the `mclust` R package (Scrucca *et al.* 2016). Medaka longevity ranges from one year in the wild in Japan to five years in the laboratory (Edeline et al. 2016 and references therein). Therefore, we allowed for one to five Gaussian component models and further allowed for different variances between the Gaussian components. Then, we selected the optimal model and corresponding number of Gaussian components according to Bayesian Information Criterion, as returned by the `mclustBIC` function (the optimal model was that yielding the highest BIC value).

We estimated medaka population dynamics in ponds through their stock-recruitment relationship, where stock in the total number of fish in March (i.e., number of age 0+ and 1+ individuals) and recruitment is the number of age 0+ individuals in Autumn of the same year. To estimate recruitment, we fitted a mixture of two Gaussian distributions to individual standard body lengths $Sdl$ :

$$
\begin{aligned}
& Sdl_i \sim \sum_{j=1}^{J} \sum_{k=1}^{K} \pi_{j,k} N\left(\mu_{j,k}, \sigma_j^2\right) \\
& \mu_{2,k} \sim N\left(\mu_{H[k]}, \sigma^2\right) \\
& \mu_{1,k} = \delta_k \mu_{2,k} \\
& \delta_k \sim U(0,1)
\end{aligned} \qquad \text{(1a)},
$$

where $i$ indexes individuals ($n$ = 17908), $j$ indexes age groups (age 0+ *vs*. 1+ such that $J$ = 2), $k$ indexes a sampling event, i.e., indexes one population in a particular year and month ($K$ = 109 sampling events), $N$ is the normal distribution, and $U$ is the uniform distribution. $H[k]$ indexes the harvest treatment (harvested *vs*. non harvested) associated with sampling event $k$. $\pi_{j,k}$ is the proportion of age $j$ individuals at each sampling event $k$ such as for each $k$:

$$\pi_j \geq 0, \sum_{j=1}^{J} \pi_j = 1 \qquad \text{(1b)}.$$

Indexes in line 1 in Eq. 1a show that our model estimated a mean standard body length separately for each age group at each sampling event, while body length variance was assumed to vary only with age. Line 2 in Eq. 1a shows that we assumed the mean standard body length at age 1+ at each sampling event, $\mu_{2,k}$, to be a normally-distributed random variable with higher-hierarchical mean specific to each harvest treatment, because harvesting was expected to restrict the maximum body size of medaka. Lines 3-4 in Eq. 1a show that mean standard body length of age 0+ medaka at each sampling event, $\mu_{1,k}$, was estimated as proportional to $\mu_{2,k}$ with a proportionality constant $\delta_k$ following a uniform distribution between 0 and 1. Model 1 provided us with MCMC (see below) age samples for each individual fish in the dataset, allowing us to compute age-specific exploitation rates by the fishery that were on average 58% (95% credible interval 34-72 %) on age 0+ juveniles, and 96 % (95% credible interval 92-98 %) on age 1+ adults.

Model (1) above allowed us to estimate the absolute number $R_k$ of age 0+ medaka recruits at each November sampling event $k$ ($n$ = 60 November sampling events). We then visualized the strength of negative density-dependence in pond medaka populations by plotting (Fig. 3b in the main text) Ricker "stock-recruitment" relationships between $R_k$ and the number $S_k$ of fish released in March (stock of spawners):

$$\begin{aligned} & R_k \sim P(\lambda_k) \\ & \ln(\lambda_k) = \ln(S_k) + \alpha_{Year[k]} + \beta_{Year[k]} S_k \end{aligned} \quad (3),$$

where $P$ is the Poisson distribution and $Year[k]$ indexes indicate that one Ricker curve was fitted for each year from 2012 to 2016.

*Larvae counts*

Larvae counts $L$ were highly overdispersed and followed a zero-inflated negative binomial distribution, which we modelled as (Ntzoufras 2009):

$$\begin{aligned} & L_i \sim NB(\phi_i, r_{H[i]}) \\ & \phi_i = \frac{r_{H[i]}}{r_{H[i]} + \lambda_i (1 - \theta_i)} \\ & \ln(\lambda_i) = \alpha_{k[i]} + \beta_{H[i]} + \gamma_{H[i]} Day_i + \delta_{H[i]} Day_i^2 \\ & \alpha_{k[i]} \sim N(0, \sigma_\alpha^2) \end{aligned} \quad (4a),$$

where $NB$ is the negative binomial distribution with success probability $\phi$ and number of failures $r$, and subscript $i$ indexes counts from a given observer in a given population on a given sampling day ($n$ = 2004 counts). Lines 3 and 4 in Eq. (4a) show that we modelled positive (non-zero) counts $\lambda$

as a harvest treatment-specific, 2nd order polynomial of the day of year (scaled to 0 mean), with a normally-distributed random effect of $k$ corresponding, as above, to a given population in a given year ($n$ = 36).

The $\theta$ latent variable for absence of larvae was modelled as a Bernoulli process being a 2nd order polynomial of the day of year :

$$\theta_i \sim B(\psi_i)$$
$$\ln\left(\frac{\psi_i}{1-\psi_i}\right) = \epsilon + \zeta \, Day_i + \eta \, Day_i^2 \quad \text{(4b),}$$

where $B$ is the Bernoulli distribution with probability of larvae absence $\psi$ .

Line 2 in Eq. 4a shows that we allowed for the $r$ parameter, which enters in the computation of the variance of the distribution (Ntzoufras 2009), to be different among the two harvest treatments $H$ . Harvest treatment-specific mean larvae count is given by $E(L_H) = \bar{\lambda}_H (1-\bar{\theta})$ and variance by $var(L_H) = \bar{\lambda}_H (1-\bar{\theta})(\bar{\lambda}_H (1-\bar{\theta}) + r_H)$ . In Table S1, we computed the dispersion index in each harvest treatment as $DI_H = E(L_H) / var(L_H)$ (Ntzoufras 2009).

*Zooplankton and filamentous algae in ponds*

We estimated the effect of medaka fishing on zooplankton abundances ($n$ = 960 observations) using a zero-inflated negative binomial GLMM (e.g., model 4 described above). The linear predictor was the same for both positive counts and the latent variable for absence, and included as fixed effects a medaka fishing-by-zooplankton taxon interaction ($n$ = 2*6 = 12 levels), and as normally-distributed random intercepts the pond ($n$ = 12 levels), sampling date ($n$ = 11 levels), and enumeration method ($n$ = 3 levels). We estimated an effect of medaka fishing on % of pond covered by filamentous algae ($n$ =

234 observations) using a negative binomial GLMM that included as fixed effect medaka fishing (*n* = 2 levels) and as random intercepts the pond (*n* = 12 levels), sampling date (*n* = 7 levels) and the observer (*n* = 9 levels).

*Somatic growth rates and trajectories of $F_1$ progeny in the laboratory*

We estimated harvest-by-food interactions on medaka growth trajectories using a 2nd order polynomial regression of standard body length $Sdl$ on age (measured in days-post-hatch):

$$
\begin{aligned}
& Sdl_i \sim N(\mu_i, \sigma_i^2) \\
& \mu_i = \alpha_{P[i]} + \beta_{H[i]} + (\gamma_{H[i],F[i]} + \delta_{P[i]}) * Age_i + \eta\, Age_i^2 \\
& \alpha_{P[i]} \sim N(0, \sigma_\alpha^2) \\
& \delta_{P[i]} \sim N(0, \sigma_\delta^2) \\
& \ln(\sigma_i^2) = \mathrm{A}_{H[i],F[i]} + \mathrm{B}_{H[i],F[i]}\, Age_i
\end{aligned} \qquad (5),
$$

where $i$ indexes length observations (*n* = 1144 observations from 104 individuals), $H[i]$ indexes the harvest treatment associated with observation $i$ (*n* = 2 levels), $H[i],F[i]$ indexes the interaction of harvest treatment and food environment (*n* = 2 * 3 = 6 levels), and $P[i]$ indexes the parental breeding pair associated with observation $i$ (*n* = 36 pairs), treated as a normally-distributed random effect on both size-at-hatch $\alpha$ and the linear somatic growth rate $\gamma$ . The six $\gamma$ parameters in Eq. 5 estimate the slope of the age effect on $Sdl$ and provided somatic growth rates as plotted in Fig. 5a of the main text. The random pair effects on somatic growth rate, $\delta_P$ , are shown in Fig. S2.

In this model, we assumed both linear somatic growth rate $\gamma$ and the regression of (ln-transformed) residuals variance on age to be different among harvest treatments and food environments (lines 2 and

5 in Eq. 5, respectively). In contrast, size-at-hatch $\beta_{H[i]}$ was allowed to vary only due to harvest treatment because food environments were applied only starting from 15 dph.

*Probabilistic maturation reaction norms of $F_1$ progeny in the laboratory*

Probabilistic maturation reaction norms (PMRNs) describe the probability that an immature individual at a given age and size will mature during a given interval of time (Heino *et al.* 2002). Provided that plasticity in the maturation process is captured by growth trajectories, PMRNs separate the effects of evolution from plasticity on maturation. PMRNs have been extensively used to explore genetic effects of exploitation on the maturation process in wild populations (Olsen *et al.* 2004; Heino & Dieckmann 2008). We fitted a Bernouilli model to individual medaka maturity (0 or 1) data $y_i$ , truncated so as to keep only the first maturity event for each individual (Heino & Dieckmann 2008):

$$\begin{aligned} & y_i \sim B(\mathrm{M}_i) \\ & \ln\left(\frac{\mathrm{M}_i}{1-\mathrm{M}_i}\right) = \alpha_{P[i]} + \beta_{H[i]} + \gamma_{H[i]} Age_i + \delta_{H[i]} Sdl_i \\ & \alpha_{P[i]} \sim N(0, \sigma_\alpha^2) \end{aligned} \qquad (6),$$

where $\mathrm{M}$ is maturity probability. Other subscripts or variables are as described in Eq. 5. The random pair effects on maturation probability at an average age and length, $\alpha_P$ , shown in Fig. S2.

*Maturation rates of $F_1$ progeny in the laboratory*

Technically, the PMRN approach assumes that observations are made at regular time intervals and, biologically, PMRNs assume that maturation is a discrete event. In the reality, however, observations are often made at irregular intervals (e.g., we observed medaka at intervals ranging from 6 to 17 days, 10 days on average), and maturation is often the threshold phenotypic expression of a continuous physiological process (Harney et al. 2013 and references therein). To bypass these problems, maturation rate models were developed that are not sensitive to the periodicity of observations and can more finely capture the physiological dynamics that underlie maturation (Van Dooren *et al.* 2005). Harney et al. (2013) have shown that maturation rate models may be approximated by fitting maturation data to standard GLMs:

$$\begin{aligned} &y_i \sim B(\mathrm{M}_i) \\ &\ln\left(\frac{\mathrm{M}_i}{1-\mathrm{M}_i}\right) = \ln(\Delta_i) + \alpha_{P[i]} + \beta_{H[i],F[i]} + \gamma\, Age_i + \delta\, Sdl_i \\ &\alpha_{P[i]} \sim N(0, \sigma_\alpha^2) \end{aligned} \quad (7),$$

where subscripts are similar to that described in Eq. 6, and the duration interval (days) between two observations $\Delta$ is included as an offset term. We used this approach to estimate harvest-by-food interaction on medaka maturation rates, in complement with the PMRN approach described above. The parameter $\beta$ captures maturation rates (in logit of maturation probability day$^{-1}$), as plotted in Fig. 5b in the main text.

*Predatory behaviour of $F_1$ progeny in the laboratory*

Counts $C_i$ of number of prey eaten by individual medaka followed a zero-inflated negative binomial distribution and were modelled similarly to larvae counts in model 4 above:

$$\begin{aligned} &C_i \sim NB(\phi_i, r_{H[i],F[i]}) \\ &\phi_i = \frac{r_{H[i],F[i]}}{r_{H[i],F[i]} + \lambda_i (1-\theta_i)} \\ &\ln(\lambda_i) = \alpha_{I[i]} + \beta_{H[i],F[i]} \\ &\alpha_{I[i]} \sim N(0, \sigma_\alpha^2) \end{aligned} \quad (8a),$$

where number of failures $r$ and positive (non-zero) counts $\lambda$ were both modelled as being different among harvest treatments $H$ in each food environment $F$, while $\alpha_{I[i]}$ was a normally-distributed random individual effect on $\lambda$ (n = 104 individuals). The $\theta$ latent variable was modelled as:

$$\begin{gathered}\theta_i \sim B(\psi_i)\\ \ln\left(\frac{\psi_i}{1-\psi_i}\right)=\gamma+\delta_{I[i]}\\ \delta_{I[i]} \sim N(0,\sigma_\delta^2)\end{gathered} \quad \text{(8b),}$$

where $\delta_I$ is a normally-distributed random individual effect.

*Analysis of deviance*

We tested for the overall statistical significance of harvest-by-food interactions on somatic growth and maturation in the laboratory using analyses of deviance. Specifically, we fitted the following models:

$$\begin{gathered}Sdl_i \sim N(\mu_i, \sigma^2)\\ \mu_i=\alpha_{H[i]}+(\beta_{H[i]}+\gamma_{F[i]}+\delta_{H[i],F[i]})Age_i+\epsilon\, Age_i^2\end{gathered} \quad \text{(9), and}$$

$$\begin{gathered}y_i \sim B(\mathrm{M}_i)\\ \ln\left(\frac{\mathrm{M}_i}{1-\mathrm{M}_i}\right)=\alpha_{H[i]}+\beta_{F[i]}+\gamma_{H[i],F[i]}+(\delta_{H[i]}+\epsilon_{F[i]}+\zeta_{H[i],F[i]})Age_i+(\eta_{H[i]}+\theta_{F[i]}+\iota_{H[i],F[i]})Sdl_i\end{gathered} \quad \text{(10),}$$

where variables and indexes are as in models (5) and (6).

*Parameter estimation*

Models 3, 9 and 10 were fitted using maximum likelihood (`glm` function, "`quasibinomial`" distribution for Eq. 10) in R 3.6.1 (R Core Team 2019). Analysis of deviance for models 9 and 10 was performed with the `anova` function using an F test to evaluate the significance of each predictor separately (Table S3). Models for the abundance of zooplankton and for % of pond covered by filamentous algae were fitted by maximum likelihood using the `glmmTMB` library of the R software (Brooks *et al.* 2017). Other models were fitted by Markov chain Monte Carlo (MCMC) in JAGS 4.2.0 (Plummer 2003) through the jagsUI package (Kellner 2019). To ease model convergence and avoid slope-intercept correlations, all numerical predictors were scaled to zero mean and, in case of Bernoulli distributions with logit links, further standardized to 0.5 standard deviation (Gelman *et al.* 2008). For each model, we ran three independent MCMC chains thinned at a period of 5 iterations until parameter convergence was reached, as assessed using the Gelman–Rubin statistic (Gelman & Rubin 1992).

Parameter estimates for models 4-6 and 8 are provided in Table S2. Statistical significance of harvest- and food-treatment effects reported in the main text was assessed from the posterior distributions of parameter differences in a test equivalent to a bilateral *t* test. In these tests, the MCMC P-value was twice the proportion of the posterior for which the sign was opposite to that of the mean posterior value. Priors were chosen to be weakly informative. In model 1 we used a Dirichlet prior for $\pi_{j,k}$ and prevented label switching by assigning age class 0+ to fish shorter than 8 mm and age class 1+ and older to fish longer than 35 mm (Chung *et al.* 2004).

We assessed goodness of fit of our models by using a Bayesian P-value (Gelman *et al.* 1996). Briefly, we computed residuals for the actual data as well as for synthetic data simulated from estimated model parameters (i.e., residuals from fitting the model to ‘‘ideal’’ data). The Bayesian P-value is the proportion of simulations in which ideal residuals are larger than true residuals. If the model fits the data well, the Bayesian P-value is close to 0.5. Bayesian P values for our models ranged from 0.47 to 0.57 and were on average 0.51, indicating excellent model fit to the data.

### APPENDIX II. NATURAL SELECTION ON BODY SIZE IN MEDAKA

We suggest that natural selection favoured small-bodied medaka in the wild, but large-bodied medaka in ponds. In the wild, medaka starve to death during their first reproductive bout while reaching age 1+, suggesting that small-bodied juvenile medaka exclude their large-bodied parents in exploitative competition for food (Edeline et al. 2016 and references therein). This is presumably because the complex habitat structure and relatively low population densities that prevail in the wild reduce interference and make competition to operate mainly through food exploitation, in which case a small body size provides fish with a strong competitive advantage (Persson *et al.* 1998; Persson & De Roos 2006). This natural selection regime in the wild was shifted in our experimental ponds, where overcompensating stock-recruitment curves mediated by increased juvenile mortality demonstrate that large-bodied adults dominated small-bodied juveniles. Compared to the wild, ponds had drastically reduced habitat complexity and probably also higher population densities. These environmental changes likely shifted competition to operate mainly through interference, which was shown to favour larger body sizes in multiple systems (Post *et al.* 1999; Calsbeek & Smith 2007; Reichstein *et al.* 2013; Le Bourlot *et al.* 2014). In fish, interference is often associated with cannibalism which also favours larger body sizes (Claessen *et al.* 2000, 2004).

### APPENDIX III: SUPPLEMENTARY RESULTS

**Table. S1. Inference of number of age classes in pond medaka populations from body length distributions using model-based clustering.** Models including one to five Gaussian components were fitted to body-lengths separately for each year and harvest treatment. The optimal number of Gaussian components was that corresponding to the model returning the highest BIC (Scrucca *et al.* 2016).

| Harvest treatment | Year | Optimal number of Gaussian components |
|---|---|---|
| Harvested | 2012 | 2 |
| | 2013 | 2 |
| | 2014 | 2 |
| | 2015 | 1 |
| | 2016 | 2 |
| Unharvested | 2012 | 2 |
| | 2013 | 2 |
| | 2014 | 2 |
| | 2015 | 2 |
| | 2016 | 2 |

**Table S2. Structure and MCMC parameter estimates for models 4-6 and 8.** The MCMC P-value is twice the proportion of the posterior for which the sign was opposite to that of the mean posterior value. MCMC P-values are not relevant for variance parameters that are constrained to be non-zero.

| Response | N | Distribution | Link | Effect | Mean estimate | SD of the estimate | MCMC P-value |
|---|---|---|---|---|---|---|---|
| Larvae count | 2004 | Bernoulli in ZINB | logit | Int. | -8.254 | 1.385 | 0.000 |
| | | | | Slope of day | -6.703 | 1.008 | 0.000 |
| | | | | Slope of day squared | 10.356 | 1.800 | 0.000 |
| | | Negative binomial in ZINB | ln | Int. no-harvest | 2.167 | 0.250 | 0.000 |
| | | | | Int. harvest | 1.250 | 0.251 | 0.000 |
| | | | | Slope of day no-harvest | 0.292 | 0.080 | 0.000 |
| | | | | Slope of day harvest | 2.211 | 0.141 | 0.000 |
| | | | | Slope of day squared no-harvest | -0.472 | 0.127 | 0.000 |
| | | | | Slope of day squared harvest | -1.638 | 0.182 | 0.000 |
| | | | | Dispersion index no-harvest | 5.372 | 1.125 | |
| | | | | Dispersion index harvest | 3.996 | 0.780 | |
| | | | | SD of year by pond effect (random) | 0.977 | 0.133 | |
| Standard body length | 1144 | Gaussian | Identity | Int. no-harvest | 4.410 | 0.106 | 0.000 |
| | | | | Int. harvest | 4.548 | 0.099 | 0.000 |
| | | | | Slope of age no-harvest low food | 0.224 | 0.005 | 0.000 |
| | | | | Slope of age harvest low food | 0.210 | 0.005 | 0.000 |
| | | | | Slope of age no-harvest medium food | 0.250 | 0.005 | 0.000 |
| | | | | Slope of age harvest medium food | 0.231 | 0.005 | 0.000 |
| | | | | Slope of age no-harvest high food | 0.263 | 0.005 | 0.000 |
| | | | | Slope of age harvest high food | 0.248 | 0.004 | 0.000 |
| | | | | Slope of age squared | -0.001 | 0.000 | 0.000 |
| | | | | Int. residual variance no-harvest low food | -0.012 | 0.149 | 0.921 |
| | | | | Int. residual variance harvest low food | -0.547 | 0.123 | 0.000 |
| | | | | Int. residual variance no-harvest medium food | -0.605 | 0.149 | 0.000 |
| | | | | Int. residual variance harvest medium food | -0.377 | 0.138 | 0.007 |
| | | | | Int. residual variance no-harvest high food | -0.534 | 0.130 | 0.001 |
| | | | | Int. residual variance harvest high food | -0.296 | 0.151 | 0.063 |
| | | | | Slope of age residual variance no-harvest low food | -0.005 | 0.003 | 0.063 |
| | | | | Slope of age residual variance harvest low food | 0.011 | 0.002 | 0.000 |
| | | | | Slope of age residual variance no-harvest medium food | 0.000 | 0.003 | 0.985 |
| | | | | Slope of age residual variance harvest medium food | 0.010 | 0.002 | 0.000 |
| | | | | Slope of age residual variance no-harvest high food | 0.011 | 0.002 | 0.000 |
| | | | | Slope of age residual variance harvest high food | -0.011 | 0.003 | 0.001 |
| | | | | SD of parental pair effect on int. (random) | 0.014 | 0.002 | |
| | | | | SD of parental pair on slope of Age effect (random) | 0.303 | 0.069 | |

**Continues on the next page.**

**Table S2 continued.**

| Response | N | Distribution | Link | Effect | Mean estimate | SD of the estimate | MCMC P-value |
|---|---|---|---|---|---|---|---|
| Maturation probability | 591 | Bernoulli | logit | Int. no-harvest low food | -8.372 | 0.922 | 0.000 |
| | | | | Int. harvest low food | -6.477 | 0.723 | 0.000 |
| | | | | Int. no-harvest medium food | -6.009 | 0.685 | 0.000 |
| | | | | Int. harvest medium food | -5.905 | 0.626 | 0.000 |
| | | | | Int. no-harvest high food | -6.063 | 0.735 | 0.000 |
| | | | | Int. harvest high food | -6.544 | 0.676 | 0.000 |
| | | | | Slope of age | 1.554 | 0.908 | 0.090 |
| | | | | Slope of length | 7.247 | 1.164 | 0.000 |
| | | | | SD of parental pair effect on int. (random) | 1.469 | 0.355 | |
| Prey count | 311 | Bernoulli in ZINB | logit | Int. | -1.960 | 0.541 | 0.000 |
| | | | | SD of individual effect (random) | 0.903 | 0.534 | |
| | | Negative binomial in ZINB | ln | Int. no-harvest, low food | 2.035 | 0.208 | 0.000 |
| | | | | Int. harvest, low food | 1.848 | 0.231 | 0.000 |
| | | | | Int. no-harvest, medium food | 1.928 | 0.245 | 0.000 |
| | | | | Int. harvest, medium food | 0.986 | 0.286 | 0.001 |
| | | | | Int. no-harvest, high food | 0.357 | 0.270 | 0.188 |
| | | | | Int. harvest, high food | 0.672 | 0.309 | 0.025 |
| | | | | Dispersion index no-harvest, low food | 2.388 | 0.722 | |
| | | | | Dispersion index harvest, low food | 5.994 | 2.141 | |
| | | | | Dispersion index no-harvest, medium food | 6.509 | 3.857 | |
| | | | | Dispersion index harvest, medium food | 5.012 | 2.357 | |
| | | | | Dispersion index no-harvest, high food | 2.033 | 0.710 | |
| | | | | Dispersion index harvest, high food | 5.708 | 2.642 | |
| | | | | SD of individual effect (random) | 0.681 | 0.136 | |

**Table S3. Effect of medaka fishing on medaka food in ponds.** Zooplankton abundances are counts per liter and abundances of filamentous algae are % of pond surface covered. Predictions were obtained from statistical models described in the SI Appendix I. There was a large variability in zooplankton counts due to the effects of the pond, sampling date and enumeration method, and the positive effect of medaka fishing was statistically significant on *Asplanchna* sp. (probability of presence, $p = 0.033$), copepodites of calanoids (non-zero abundances, $p < 0.001$) and Cladocerans (non-zero abundances, $p < 0.001$) before June but not after (results not shown), probably because of medaka recruitment that increased medaka density in unharvested ponds. The effect of medaka fishing on filamentous algae was statistically significant ($p < 0.002$).

| **Taxa** | **Medaka treatment** | **Predicted Count / L (zooplankton) or % pond surface cover (filamentous algae)** | |
|---|---|---|---|
| | | **Mean** | **SE** |
| *Asplanchna* sp. | Harvested | 14.2 | 15.5 |
| | Unharvested | 8.5 | 9.6 |
| Copepodites of calanoids | Harvested | 161.6 | 150.0 |
| | Unharvested | 85.2 | 79.1 |
| Cladocerans | Harvested | 51.4 | 48.1 |
| | Unharvested | 18.1 | 17.1 |
| Copepodites of cyclopoids | Harvested | 2.7 | 2.6 |
| | Unharvested | 3.2 | 3.1 |
| Copepod *nauplii* | Harvested | 213.3 | 197.6 |
| | Unharvested | 215.2 | 199.8 |
| Small rotifers | Harvested | 7018.9 | 6537.0 |
| | Unharvested | 6939.2 | 6458.5 |
| Filamentous algae | Harvested | 18.5 | 11.5 |
| | Unharvested | 1.3 | 0.8 |

**Table S4. Analysis of deviance for GLMs testing for the harvest-by-food interaction on life-history traits in laboratory-born $F_1$ medaka progeny.** The “Deviance” column gives the reduction in the residual deviance as each predictor is added in turn into the model. The P-values compare the reduction in deviance to the residual deviance in an F test.

| Trait | Distribution | Link | Predictor | Df | Deviance | Resid. DF | Resid. Dev | F | P-val |
|---|---|---|---|---|---|---|---|---|---|
| Body length | Gaussian | Identity | Harvesting | 1 | 163 | 1130 | 20588 | 155 | **<0.0001** |
| | | | Food | 2 | 658 | 1128 | 19929 | 312 | **<0.0001** |
| | | | Age | 1 | 18086 | 1127 | 1843 | 17124 | **<0.0001** |
| | | | Age^2 | 1 | 469 | 1126 | 1374 | 444 | **<0.0001** |
| | | | Harvesting x Food | 2 | 2 | 1124 | 1372 | 1 | 0.3282 |
| | | | Harvesting x Age | 1 | 22 | 1123 | 1350 | 21 | **<0.0001** |
| | | | Food x Age | 2 | 167 | 1121 | 1183 | 79 | **<0.0001** |
| | | | Harvesting x Food x Age | 2 | 1 | 1119 | 1182 | 0 | 0.7160 |
| Maturation | Bernoulli | Logit | Harvesting | 1 | 0 | 589 | 528 | 0 | 0.5758 |
| | | | Food | 2 | 6 | 587 | 522 | 6 | **0.0041** |
| | | | Age | 1 | 135 | 586 | 387 | 253 | **<0.0001** |
| | | | Length | 1 | 57 | 585 | 329 | 107 | **<0.0001** |
| | | | Harvesting x Food | 2 | 7 | 583 | 323 | 6 | **0.0022** |
| | | | Harvesting x Age | 1 | 3 | 582 | 320 | 5 | **0.0195** |
| | | | Food x Age | 2 | 11 | 580 | 309 | 11 | **<0.0001** |
| | | | Harvesting x Length | 1 | 0 | 579 | 308 | 0 | 0.5630 |
| | | | Food x Length | 2 | 2 | 577 | 307 | 2 | 0.2142 |
| | | | Harvesting x Food x Age | 2 | 15 | 575 | 292 | 14 | **<0.0001** |
| | | | Harvesting x Food x Length | 2 | 1 | 573 | 290 | 1 | 0.3197 |

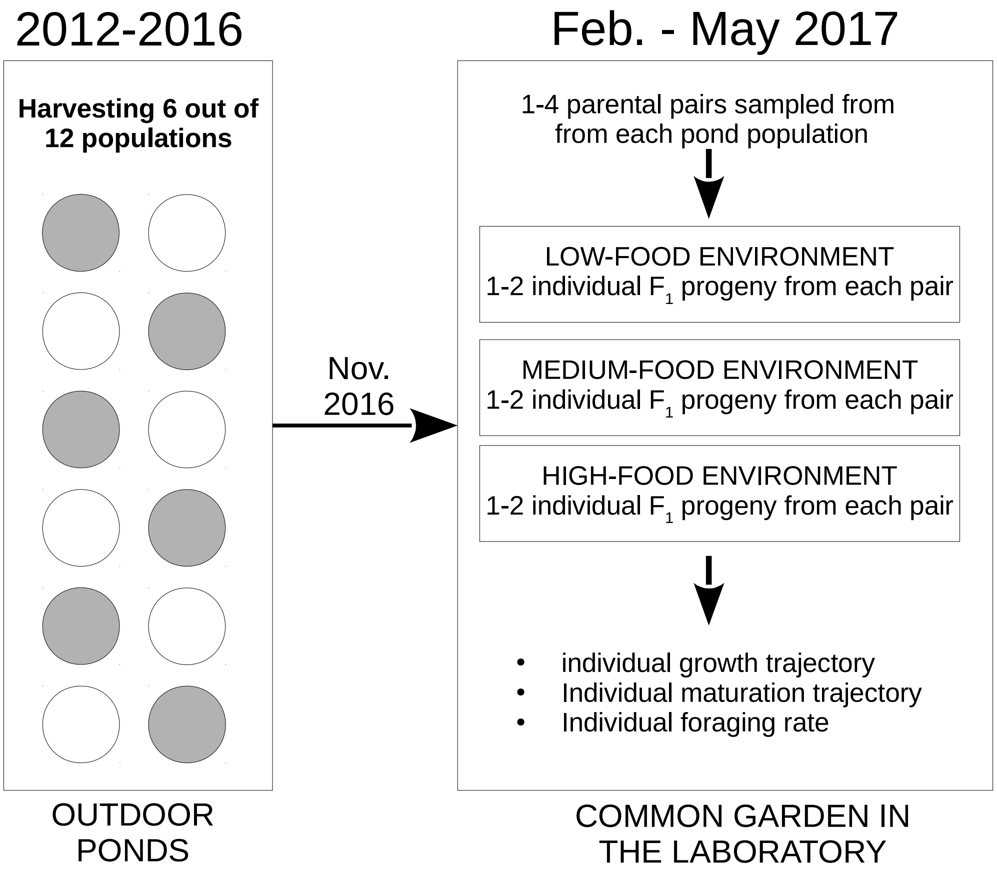


**Fig. S1. Experimental design.** In 2012, 12 independent populations of medaka were introduced in 12, 10 m$^2$ outdoor ponds and maintained naturally with no added food. Each year from 2012 to 2016, the 12 populations were sampled (98% catch rate) in March and November, and each fish was individually weighed. Each year in March in six populations (shaded), only the 19% smallest-bodied individuals from the catch were released, while all individuals were released in the other six populations (unshaded). Each year in November, all fish were released after weighing, except in November 2016 when a random sample of 6-10 fish (mean 9.6) from each population was kept and transferred to the laboratory to serve as parents in a common garden experiment. In 2017, parents originating from the same population were mated and their progeny was distributed in individual tanks under three food environments (Low, Medium, High), where we measured their individual somatic growth rate, maturation trajectory, and foraging rate.

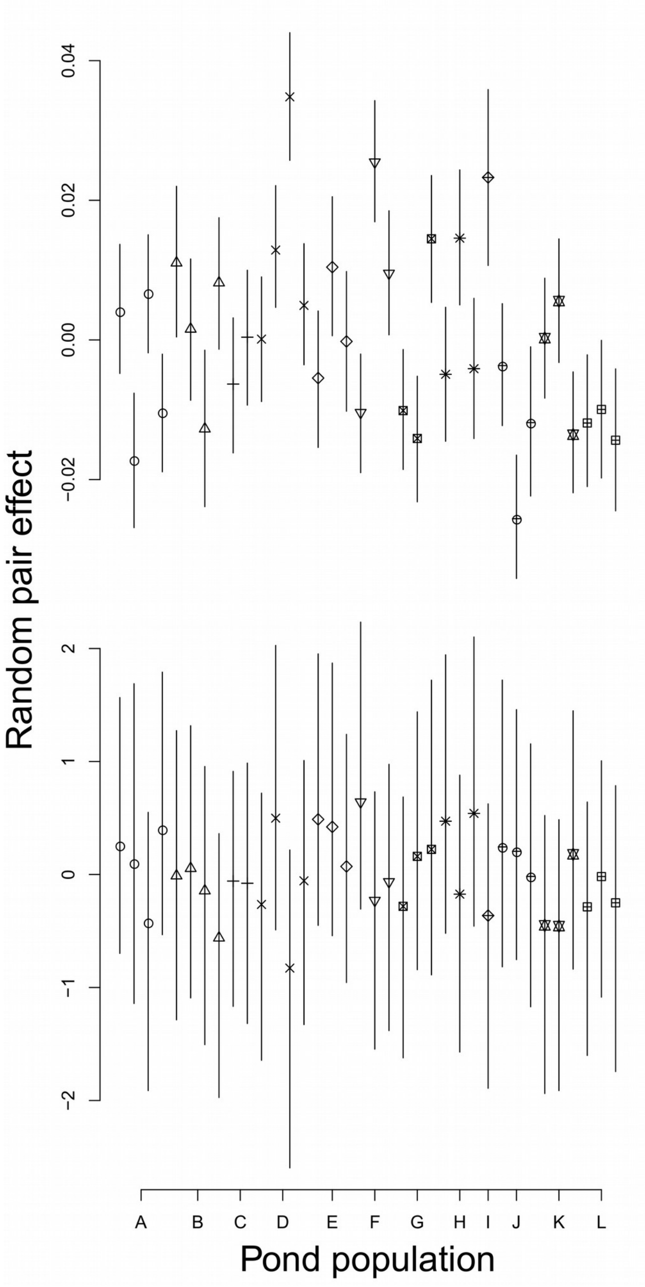


**Fig. S2. Random effects of breeding pairs on the somatic growth rate (top) and maturation probability (bottom) of $F_1$ medaka progeny in the laboratory.** Effects were estimated by MCMC from models 5 and 6, as described above. Points show median MCMC estimates with 95% credible intervals. Effects for somatic growth rate are in mm $day^{-1}$ and effects on maturation are in logit (probability). Symbols correspond to the pond population of origin (coded on the x axis from A to L). Populations A, D, F, G, J and K were harvested while other populations were unharvested.

368
